# Earlier and More Robust Sensorimotor Discrimination of ASL Signs in Deaf Signers During Imitation

**DOI:** 10.1101/2020.01.31.929208

**Authors:** Lorna C. Quandt, A. S. Willis

## Abstract

**Background:** Prior research suggests that the amount of experience an individual has with an action influences the degree to which the sensorimotor systems of their brain are involved in the subsequent perception of those actions. Less is known about how action experience and conceptual understanding impact sensorimotor involvement during imitation. We sought to explore this question by comparing a group of sign language users to a group of non-signers. We pitted the following two hypotheses against each other: 1) Deaf signers will show increased sensorimotor activity during sign imitation, and greater differentiation between sign types, due to greater prior experience and conceptual understanding of the signs; versus 2): Deaf signers will show less sensorimotor system activity and less differentiation of sign types in the sensorimotor system, because for those individuals sign imitation involves language systems of the brain more robustly than sensorimotor systems. We collected electroencephalograms (EEG) while the two groups imitated videos showing one-handed and two-handed ASL signs. Time-frequency data analysis was performed on alpha- and beta-range oscillations while they watched signs with the intent to imitate, and imitated the signs. During observation, deaf signers showed early differentiation in alpha/beta power between the one- and two-handed sign conditions, whereas hearing non-signers did not discriminate between the sign categories this way. Significant differences between groups were seen during sign imitation, wherein deaf signers showed desynchronization of alpha/beta EEG signals, and hearing non-signers showed increased power. The study suggests that in an imitative context, deaf signers engage anticipatory motor preparation in advance of action production, while hearing non-signers engage slower, more memory-related processes to help them complete with the complex task.

## Introduction

To the unlearnt eyes, the blur of sign language is challenging to follow. Fluent deaf sign language users produce and understand layers of serial and simultaneous actions with their body, hands, and face, easily communicating complex linguistic content through the visual-manual modality. Recent research has shed light on how signers exhibit differences in visual perception and motion processing, in some cases in response to sign-related stimuli (Kubicek & Quandt, 2019), and in other cases in response to non-sign stimuli (Peressotti, Scaltritti, & Miozzo, 2018; Quandt & Kubicek, 2018; Williams, Darcy, & Newman, 2016). However, little is known about how people perceive signs in imitative contexts. In an imitative context, an observer watches with the intent to reproduce the action that they see. Many cognitive neuroscience studies of signed language processing engage participants in a unidirectional sign-perception task, such as lexical decision tasks, in which the participants are not intending to produce signed responses. While non-imitative tasks provide important information about how the brain processes signs, they do so in a context which is removed from sign language as a bi-directional, interactive, and communicative form of action. Here, we consider how deaf signers and hearing non-signers process signs when asked to imitate them—a task which presents a significant challenge to most hearing non-signers.

## Neural Correlates of Sign Language Perception and Production

There are clear similarities in the neural organization of the language production networks for spoken and signed languages (Emmorey, Mehta, McCullough, & Grabowski, 2016; MacSweeney, Capek, Campbell, & Woll, 2008). Mental representations and processing of signs share many correlates with the representation and processing of speech (Blanco-Elorrieta, Kastner, Emmorey, & Pylkkänen, 2018; Corina et al., 1999; Evans, Price, Diedrichsen, Gutierrez-Sigut, & MacSweeney, 2019; Petitto et al., 2016). For instance, deaf signers recruit the left superior temporal gyrus when seeing sign language phrases, much like hearing speakers do when listening to speech (MacSweeney et al., 2002). While sign languages and spoken languages have fundamental similarities in their neural organization due to the amodality of phonetic information (Emmorey, McCullough, Mehta, & Grabowski, 2014; Petitto et al., 2016), there are also important differences in the neurobiology of how spoken and signed languages are represented in the brain (Emmorey et al., 2014; Evans et al., 2019).

The long-term use of sign language seems to bring about certain changes in perception and cognition. One possible cause of the adaptive reorganization of the language processing network in signers is the demand of producing and perceiving linguistic content through several simultaneously-moving body parts. Comparing the perception of fingers, hands, and faces with the perception of lip, tongue, and vocal cord movement, there is a considerable difference in the amount of visible motion involved. Likewise, in production, sign language involves larger scale movements of the body compared to the smaller movements required for articulating speech. As such, it is not entirely surprising that long-term experience with sign language may bring about some differences in perceptual abilities related to movement. Some differences in motion-related networks between signers and non-signers suggest that these regions adapt to the perception and production demands of the signed language modality (Allen, Emmorey, Bruss, & Damasio, 2013; Emmorey et al., 2016; Kanazawa et al., 2017).

For instance, visual perceptual differences in signers are well-documented. Sign language users show differences in low level motion perception (Bosworth, Petrich & Dobkins, 2013), perception of sign-related body movements (Poizner, 1983) or positioning (Almeida, Poeppel, & Corina, 2016), visual attention, and the spatial maps of where visual attention is allocated in the focal region and/or periphery of the visual field (Dye, Baril, & Bavelier, 2007; Stoll & Dye, 2019). Signers may also show greater sensitivity to observed handshapes (Baker, Idsardi, Golinkoff, & Petitto, 2005; Morford, Grieve-Smith, MacFarlane, Staley, & Waters, 2008) although there are contradictory findings in this area (Gimeno-Martínez, Costa, & Baus, 2019). Overall, accumulating evidence shows that people with extensive sign language expertise exhibit differences in perception, but the full extent and nature of these differences is yet to be understood.

## Mirroring, Experience, and Sign

The human mirror neuron system (MNS) may play an important role in imitation and perception of human actions, and thus, has been the focus of a fair amount of inquiry regarding sign language (Corina & Knapp, 2006; Emmorey, 2014; Okada et al., 2016; Mole & Turner, 2017; Ostarek & Huettig, 2019). Typically, brain regions making up the MNS are recruited both during the perception and the production of actions, causing similar neural activity during observation and execution of the same action (Rizzolatti & Fabbri-Destro, 2009). The MNS appears to also be involved in response to aural input (Jenson et al., 2014; Saltuklaroglu, Harkrider, Thornton, Jenson, & Kittilstved, 2017; Thornton, Harkrider, Jenson, & Saltuklaroglu, 2017), perception of speech reading (Swaminathan et al., 2013), and perception of robotic voices (Di Cesare, Errante, Marchi, & Cuccio, 2017). Research supports the notion that the MNS is somehow involved in language processing, although there are significant disagreements over its role (Emmorey, 2014; Gallese, Gernsbacher, Heyes, Hickok, & Iacoboni, 2011; Mole & Turner). Given that sign languages constitute a unique overlap of language, action production, and action comprehension, it is possible that unique insights may be gained from studying mirroring-related processes in groups of experienced sign language users.

Ample evidence suggests that mirroring-like activity, such as the vicarious involvement of the somatosensory and motor cortices during action perception, may be sensitive to the observer’s own past experiences. The relation between action experience and mirroring-like activity is complex, with some research suggesting that greater experience with an action leads to increased sensorimotor involvement during subsequent perception (Calvo-Merino, Ehrenberg, Leung, & Haggard, 2010; Calvo-Merino, Glaser, Grezes, Passingham, & Haggard, 2005; Calvo-Merino, Grèzes, Glaser, Passingham, & Haggard, 2006; Cannon et al., 2014; Denis, Rowe, Williams, & Milne, 2017; Quandt & Marshall, 2014). In contrast, some findings suggest the opposite effect: that with greater action experience, the sensorimotor systems of the brain become more efficient at processing observed actions (Babiloni et al., 2010; Vogt et al., 2007) and thus show less activity. In recent years it has become clear that there is likely a non-linear relationship between action experience and involvement of the sensorimotor system during observation (Gardner, Aglinskas, & Cross, 2017a, 2017b; Gardner, Goulden, & Cross, 2015), and the nature of the relationship depends in part on the specific characteristics of the action categories in question.

Viewing sign language users as “action experts” within the domain of a sign language has yielded complex and contradictory results. Some researchers have asked whether deaf signers, due to their extensive use of the hands and body for language, may show increases in mirroring activity when perceiving sign language. However, some sign language researchers determine that these processes have little to no role in in the higher-level neural and cognitive processing of sign languages (Corina & Knapp, 2006; Emmorey, Xu, Gannon, Goldin-Meadow, & Braun, 2010; Okada et al., 2016; Rogalsky et al., 2013). Several reports found no evidence to support the involvement of mirroring during sign perception (Rogalsky et al., 2013), and in fact generally suggested that signers recruit less of the sensorimotor system during sign perception, possibly because they are relying more on linguistic processing (Möttönen, Farmer, & Watkins, 2016). Some findings suggest that deaf signers’ neural representations of action processing differ from hearing non-signers’ (Corina et al., 2007; Emmorey, McCullough, Mehta, Ponto, & Grabowski, 2011; Emmorey et al., 2010; Mole & Turner, 2017). It is possible that deaf signers’ communicative action processing is highly efficient, reducing sensorimotor system activity but not disengaging from it, due to their extensive expertise in extracting meaning from complex action (Gardner et al., 2017a, 2017b; Gardner et al., 2015). As well, the consideration of mirroring-like processes in sign language users hinges upon the definition of mirroring (Mole & Turner, 2017). Here, we turn our interest toward the broader involvement of an observer’s somatosensory and motor cortices during perception of another’s action, rather than a narrowly-defined human mirror neuron system analogous to that studied originally in macaques (Corina & Knapp, 2006).

The relationship between mirroring and experience with signed language may well be non-linear. Indeed, adult American Sign Language (ASL) learners seem to rely more on mirroring-related processes when their ASL vocabulary is weak, suggesting that at least for adult learners, reliance on mirroring-like processes may constitute a compensation in the face of weak linguistic knowledge of ASL (Williams, Darcy, & Newman, 2017). However, a recent study (Kubicek & Quandt, 2019), showed that while deaf signers show overall less involvement of the sensorimotor cortices during sign perception, the specific sensorimotor characteristics of observed signs were encoded in their sensorimotor cortices. Thus, while the overall pattern of signers showing less vicarious sensorimotor processing echoed that of prior functional neuroimaging work, both signers and non-signers were drawing upon their own sensorimotor representations to parse the details of the signs they were observing.

Analyzing the oscillatory activity of the cortex using electroencephalography (EEG) provides valuable information about what regions of the cortex are active in an action observer’s brain, which can yield fine-grained information about the timing and sensitivity of cortical sensorimotor processing during action perception and production. Common EEG measurements of sensorimotor activity in the human brain are alpha (8-13 Hz) and beta (14-30 Hz) rhythms measured at centrally-located scalp electrodes, which index the electrophysiological oscillations emanating from the primary sensory and motor cortices (Arnstein, Cui, Keysers, Maurits, & Gazzola, 2011; Bowman et al., 2017; Fox, Bakermans-Kranenburg, & Yoo, 2016; Fox, 2016). The central alpha (also termed “mu”) rhythm reflects activity in the pre-central gyrus, while the sensorimotor beta rhythm emanates from the pre-and post-central gyri (Tzagarakis, Ince, Leuthold, & Pellizzer, 2010). Sensorimotor alpha rhythms display lower power (event-related desynchronization; ERD) during action processing, whether in perception, imagination, or production, and the beta rhythm tends show similar patterns, although it may index slightly different aspects of action. The activity of these sensorimotor EEG rhythms is quite sensitive to the observer’s own experiences, with much research revealing modulations of alpha and beta power depending on the observer’s prior sensory and motor experiences with observed actions (Cannon et al., 2014; Denis et al., 2017; Quandt, Marshall, Shipley, Beilock, & Goldin-Meadow, 2012; Simonet et al., 2019). Recent work shows that sensorimotor alpha and beta rhythms can reflect mirroring-like processes during sign observation (Kubicek & Quandt, 2019) and also when deaf signers read English words (Quandt & Kubicek, 2018).

## Sensorimotor processing during sign imitation

Action perception may occur in a context where the observer is simply watching someone else’s movements, or it may occur in an imitative context, wherein the observer plans to reproduce the actions she sees. Seeing an action when one intends to copy it changes the neural profile of the observation (Decety, Chaminade, Grezes, & Meltzoff, 2002). Both imitation and action observation recruit frontal premotor, parietal, and temporo-occipital cortices (Caspers, Zilles, Laird, & Eickhoff, 2010). However, imitation particularly engages the inferior parietal cortex, primary somatosensory cortex, and the inferior frontal cortex, which is also involved in language processing (Caspers et al., 2010; Decety 2002). Many experimental tasks used in prior sign language perception studies involved passive sign observation (Corina et al., 2007; Kubicek & Quandt, 2019; MacSweeney et al., 2004; McCullough, Saygin, Korpics, & Emmorey, 2012), wherein the perceiver was not overtly planning to reproduce the signs they saw. In the current study we aimed to assess whether long-term expertise with American Sign Language (ASL) would result in enhanced or reduced involvement of the sensorimotor cortex in response to the sensorimotor characteristics of signs during imitation, in comparison to individuals unfamiliar with ASL.

We ask here how signers and non-signers perceive signs, not only looking at overall neural responses to seeing a person sign, but also the extent to which the sensorimotor systems of these observers are sensitive to the specific characteristics of the signs they see. We probed this question by asking participants to view, then imitate, signs produced either using only one hand, or signs produced using both hands. In prior work comparing neural responses to these categories of signs has yielded robust differences in sensorimotor EEG (Kubicek & Quandt, 2019; Quandt & Kubicek, 2018) and in PET measures of cortical activity (Emmorey, Mehta, McCullough, & Grabowski, 2016b). Because producing two-handed signs recruits the right sensorimotor cortex more greatly due to the involvement of the left hand, seeing the same pattern during sign observation can reveal that the observer is drawing upon his or her own internal motor plans for how the sign should be carried out (e.g., recruiting more right sensorimotor cortex when seeing a two-handed sign, which involves the left hand).

Understanding the differences in the oscillatory profile of sensorimotor EEG activity during an imitation task could improve the current understanding of how sign language users see and make sense of others’ signs. While imitation of a single sign is still far different from a natural signed conversation, the combination of sign observation with the sign production is a richer, more socially-relevant task (Krishnan-Barman, Forbes, & Hamilton, 2017). This study will examine if experience with ASL could lead to changes in sensorimotor processing of actions compared to individuals who do not know ASL.

Based on the prior work regarding these questions, we pitted the following two possible hypotheses against each other: 1) Deaf signers will show increased sensorimotor activity during sign imitation, and greater differentiation between sign types, due to greater prior experience and conceptual understanding of the signs; versus 2): Deaf signers will show less sensorimotor system activity and less differentiation of sign types in the sensorimotor system, because for those individuals sign imitation involves language systems of the brain more robustly than sensorimotor systems.

Given that recent evidence supports the latter claim (Kubicek & Quandt, 2019), we predicted our analyses would generally support that hypothesis. However, the current study uniquely engaged participants in an imitative paradigm, and given that participants were required to copy the signs they saw, we predicted that while observing with the intent to imitate, both groups would show different sensorimotor EEG responses to one-handed compared to two-handed signs, but the effect would be stronger in the Hearing Non-Signers, due to their unfamiliarity with the stimuli and the resultant need to focus on the basic physical parameters of the observed communicative actions. We also anticipated that while producing signs, Hearing Non-Signers would show more robust differentiation between one- and two-handed signs, again due to the novelty of the communicative actions requiring complex motor plans for implementation.

## Materials and Methods

### Participants

Eighteen deaf fluent signers and 19 hearing non-signers were run through the experimental protocol. Participants spanned a wide range of educational backgrounds (Table 1). Deaf fluent signers self-identified as deaf and fluent in ASL (see Table 2 for language descriptors). All participants gave their informed consent prior to the experiment and were informed of their rights in accord with the Declaration of Helsinki. An ASL version of the informed consent was shown to all deaf participants. All deaf participants were run by a native or fluent ASL signer as the lead experimenter. The study was approved by the relevant IRB and participants were paid for their time.

**Table 1.** Formal education. Self-reported highest educational degree obtained for Deaf and Hearing participants. Percentages were rounded to the nearest whole number.

|  | Deaf | Hearing |
| --- | --- | --- |
| High school | 3 (16%) | 0 |
| Some college | 2 (11%) | 1 (5%) |
| Associates | 0 | 1 (5%) |
| Bachelors | 3 (16%) | 6 (32%) |
| Some grad school | 2 (11%) | 2 (11%) |
| Masters | 8 (44%) | 8 (42%) |
| Doctorate | 0 | 1 (5%) |

**Table 2.** Demographics and ASL use. Means and standard deviations listed.

|  | Deaf | Hearing | t-test *p* value |
| --- | --- | --- | --- |
| Age (years) | 31.4 (10.7) | 27.4 (3.7) | .15 |
| Current ASL use <sup>a</sup> | 1.11 (.31) |  |  |
| ASL understanding <sup>b</sup> | 95.5 (5.1) |  |  |
| ASL production <sup>b</sup> | 92.4 (6.7) |  |  |

<sup>a</sup> Self-reported rating on a 7-point scale ranging from 1 = “All the time” to 7 = “Only on special occasions (e.g. home for the holidays)”.
<sup>b</sup> Self-rated proficiency on a sliding scale from 0 (poor) – 100 (fluent).

**Table 3.** English and ASL norms for two categories of Stimuli. 1-handed (1H) words and 2-handed (2H) words, as in Quandt & Kubicek, 2018 and Kubicek & Quandt, 2019.

|  |  | **1H words: M**<br>**(SD)** | **2H words: M**<br>**(SD)** | **t-test p**<br>**value** |
| --- | --- | --- | --- | --- |
| English norms | Word length | 5.2 (1.7) | 5.7 (2.2) | .25 |
|  | Log frequency<br>(SUBTLEX) | 5.6 (10.6) | 14.3 (43.6) | .22 |
|  | Log frequency (HAL) | 9.6 (1.7) | 10.1 (1.5) | .24 |
|  | # phonemes | 4.1 (1.1) | 4.6 (1.3) | .10 |
|  | Lexical Decision RT M | 605.6 (45.5) | 616.2 (64.4) | .40 |
|  | Lexical Decision RT SD | 202.1 (64) | 215.4 (89) | .45 |
|  | Naming RT M | 605.5 (42.6) | 607.8 (42.5) | .81 |
|  | Naming RT SD | 128 (58.1) | 130.3 (53.1) | .85 |
|  | Imageability | 509.5 (112.5) | 543.6 (79.6) | .18 |
| ASL norms | Frequency (ASL-Lex) | 4.7 (.8) | 4.6 (1.1) | .42 |
|  | Iconicity | 3.7 (1.7) | 3.8 (1.4) | .90 |
|  | Flexion | 3.4 (2.2) | 3.6 (2.2) | .69 |

### Stimuli

The ASL stimuli were video clips retrieved from ASL-LEX (Caselli, Sehyr, Cohen-Goldberg, & Emmorey, 2016). Each video clip shows a woman producing one sign. Two types of videos were used, with 40 of each of type, for a total of 80. One video type consisted of signs that only use one hand (‘one-handed’, ‘1H’; i.e. GUILT). These signs were always produced with the dominant (right) hand. The other video type consisted of signs that use two hands (‘two-handed, ‘2H’; i.e. FAMILY). Of the 40 two-handed signs, 25 were symmetrical. There were no significant differences between the signed 1-handed (1H) or 2-handed (2H) words or their English translations for any of the following measures: frequency, iconicity, flexion, phonological properties, imageability, sign onset time (ms), sign offset time (ms) and sign length (ms; for more information see Caselli et al. 2016). Action verbs and fingerspelled loan signs were not included in the stimulus set.

Each video started with a woman (a deaf native signer) sitting in a neutral position with her arms resting on her lap, and as the video continued, she raised her hands to produce an ASL sign, returning to neutral position after she finished. We conducted t-tests to assess whether the timing dynamics of one-handed and two-handed signs were similar. The onset of the sign from the start of the video clip did not differ between groups (1H: 523.0 ms, 2H: 550.6 ms, *p* = .57), nor did the offset of the signs in ms from start of video clip (1H: 1371.9 ms, 2H: 1187.9 ms, *p* = .53). The total length of the sign from onset to offset did not differ between groups (1H: 573.9 ms, 2H 637.3 ms, *p* = .22).

Using Adobe Premiere, we affixed a virtual audio trigger (a short beep) to the video clip at the onset of the sign, as defined by ASL-Lex norms (Caselli et al., 2016). The onset of the beep corresponded to the onset of the sign. The beep was never audible, but rather, was sent to a BrainProducts Trigger Box and converted into a TTL pulse. The TTL pulse was then recorded in parallel to the EEG signal in order to identify the time in the EEG signal corresponding to the time of the sign onset. Sign onset in the video was used as time 0 for all analyses.

### Experimental Procedure

After providing informed consent, participants were brought into the experimental room. While the EEG cap was prepared, they filled out a language and educational background form. They viewed the experiment in E-Prime 2.0 (Psychology Software Tools, Sharpsburg, PA) on a computer monitor ∼76 cm away. Prior to the task analyzed for this project, participants had seen the same stimuli in a different context. Participants were instructed to view each individual ASL sign, and then when prompted, to produce their own imitation of what they just saw to the best of their ability. After a brief practice session to ensure that they understood the task, the experiment started.

A total of 80 trials were presented, divided into four blocks of twenty trials each. Each sign was seen one time throughout the course of the experiment. Each trial started with a fixation point (jittered ISI = 4-5 s); then the sign clip started, with an event marker triggered at the onset of the sign (see Figure 1). After the video clip ended, a screen stating “GO” was shown for three seconds, during which the participant produced the sign they just saw. Experimental scripts are available at https://osf.io/6rtqf/.

**Figure 1.**
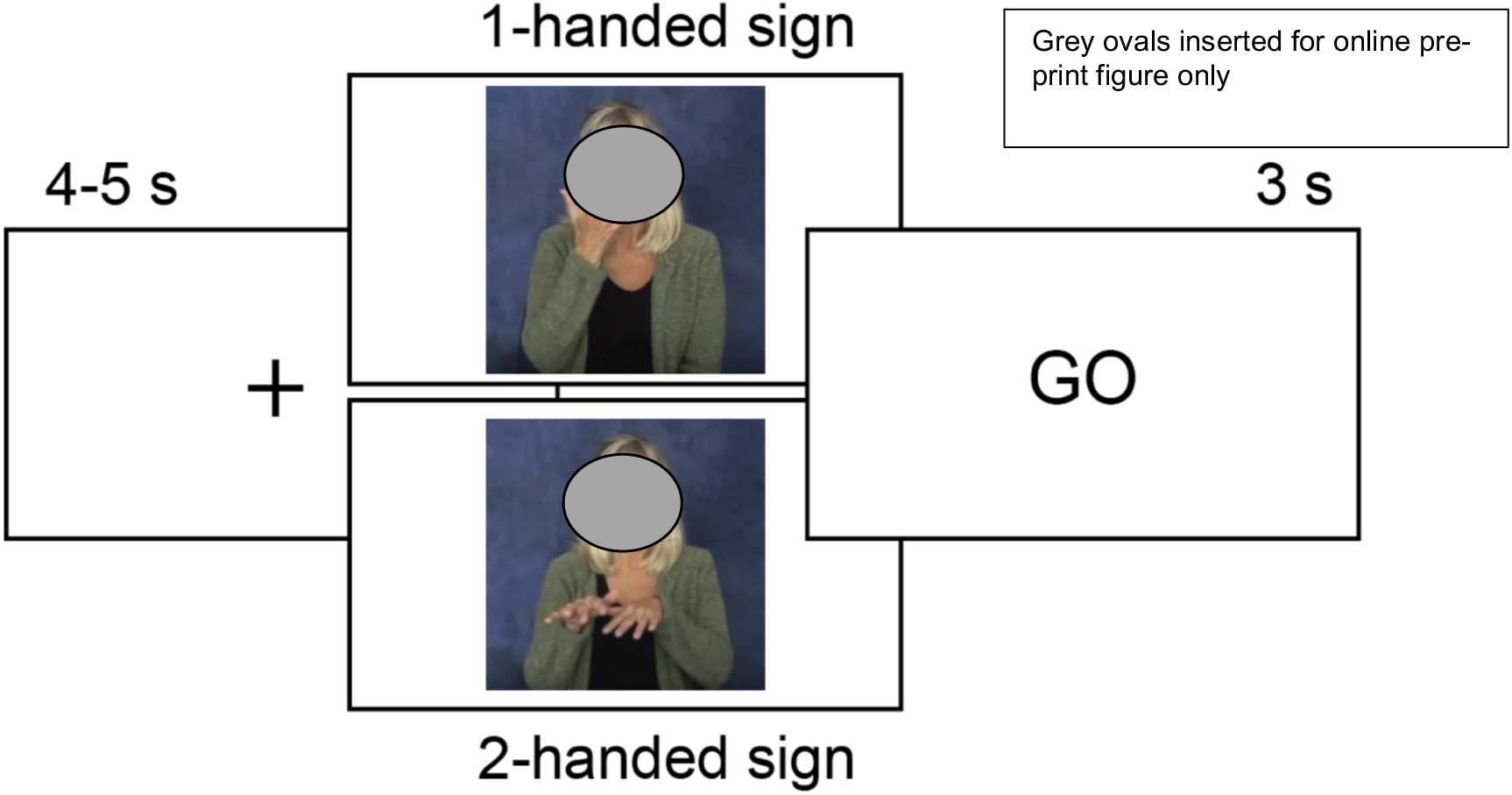
The trial structure. Participants saw one- or two-handed signs (Caselli, Sehyr, Cohen-Goldberg, & Emmorey, 2016) and then produced their own imitations of the signs when prompted by the word “GO”.

### Electroencephalographic data

EEG was recorded from 64 active Ag/AgCl electrodes using an actiCAP setup (Brain Products GmbH, Germany), with SuperVisc electrode gel. EEG data was recorded and processed at a rate of 1000 Hz. Data was recorded with a Cz reference and a grounded electrode at Afz. The EEG signals were amplified by each electrode’s active amplifier, and then by a 24-bit actiCHAMP amplifier (Brain Vision LLC, Morrisville, NC). Hardware filter settings included a high-pass filter (.53 Hz) and a low-pass filter (120 Hz).

The institutional approval for this study prohibits the public archiving of our participants’ data, however the data necessary for reproducing this work is available from the corresponding author upon request. A data-sharing agreement must be completed in advance of data sharing.

### EEG Data Processing

The EEG data were preprocessed in EEGLAB (Delorme & Makeig, 2004). Each signal was re-referenced offline to the average signal of the mastoid electrodes. The re-referenced EEG data were filtered with .01 Hz highpass and 100 Hz lowpass filters and epochs of -1500 to 5000 ms were extracted from the continuous EEG signal around the onset of the sign (time 0). Each epoch was labeled as 1H or 2H depending on condition (one-handed or two-handed). The time period of -1500 ms to -1000 was used as a baseline for all analyses. The mean signal during this time period was removed from the epochs of interest. During the baseline period there was a fixation cross visible, and for a small number of trials, the baseline included the signing model appearing on screen (this applies to clips with a longer period between clip starting and sign onset). A study file was formed with two conditions (1H signs; 2H signs) and two groups, Deaf Signers and Hearing Non-Signers. We analyzed alpha and beta range frequencies at all scalp electrodes (rather than focusing only on the mu rhythm at central electrodes) to ensure that the effects seen at central electrode sites are a result of sensorimotor activity (Arnstein et al., 2011), and not reflecting an alpha/beta-range effect present across the scalp. Here, we primarily refer to alpha and beta frequency activity, while acknowledging that the alpha-range activity occurring primarily in the central region is likely “mu” activity reflecting activity in the pre-and-post-central regions (Bowman et al., 2017).

### Statistical Analysis

#### Full scalp analysis

A 2 x 2 ANOVA with one paired condition (1H signs; 2H signs) and unpaired groups (Deaf; Hearing) was performed on two different windows within each trial. First, a window during the observation of the sign, and second, a window while participants started to produce their own imitation of the sign. This statistical comparison was performed across the full scalp to observe any significant differences in the amplitude of the frequencies of interest. We focused our analysis on four frequency bands: low alpha (8-10 Hz), high alpha (11-13 Hz), low beta (14-17 Hz), and high beta (18-25 Hz). Those frequencies were selected based on previous work analyzing sensorimotor activity during action processing (Denis et al., 2017; Quandt & Marshall, 2014; Simon & Mukamel, 2016). The epoch from 0 to 2000 ms (time 0 = onset of sign) was divided into 8 different time bins. The first four time bins (0 - 250 ms, 250 - 500 ms, 500 - 750 ms, and 750 - 1000 ms) were assessed together as part of the “observation” window, during which time the signing model was producing a sign. The last four time bins (1000 - 1250 ms, 1250 - 1500 ms, 1500 - 1750 ms, and 1750 - 2000 ms) constituted the “production” window, during which time the sign video was ending, and the participant started to produce his or her imitation. Due to the variation in the sign clip length, the production window reflects a variety of processes. We analyzed the length of the ASL videos to map out the content of the production window in the following way. The first quartile of elapsed time between sign onset and GO appearing on the screen was 1059 ms, the median elapsed time was 1251 ms, and the third quartile was 1401 ms (all times relative to sign onset, time 0). Thus, the production window used in this analysis encompassed participants’ initiation of movement, as well as some preparatory time in advance of action production.

In an effort to limit spurious findings, we decided *a priori* that a statistically significant effect would be included in our results only if it occurred at a cluster of three or more adjacent electrodes. We used an adjusted *p* value of .016 (.05 / 3) as our significance threshold for this determination to further control for multiple comparisons. This statistical approach has been used in prior work (Kubicek & Quandt, 2019; Quandt & Kubicek, 2018), and scripts are available at https://osf.io/6rtqf/.

#### Central region analysis

We performed a closer analysis at electrodes falling within our region of interest: the 21 electrodes that lay over the central region of the scalp, above the primary sensorimotor region of the brain: FC5, FC3, FC1, FCz, FC2, FC4, FC6, C5, C3, C1, Cz, C2, C4, C6, CP5, CP3, CP1, CPz, CP2, CP4, and CP6. We opted to focus analyses on these electrodes since alpha and beta rhythms at central electrodes are associated with activity in pre- and post-central gyri, which are key regions of the mirroring system (Arnstein et al., 2011; Perry & Bentin, 2009; Ritter, Moosmann, & Villringer, 2009). For each electrode, we computed time-frequency transforms from 8 - 25 Hz for observation and production windows. For these ROI analyses, the observation window extended from -750 to 1500 ms and the production window extended from 1000 - 3000 ms. The longer time windows, which included the time prior to sign onset as well as extending well into the participants’ own imitations, allowed us to more fully visualize the oscillatory dynamics unfolding over the course of the task. The same 2 x 2 ANOVA design was implemented for the ROI analyses, with one paired condition (1H signs; 2H signs) and unpaired groups (Deaf; Hearing). For these ROI analyses, a *p* value threshold of .05 was used, with a false detection rate (FDR) correction applied (Benjamini & Hochberg, 1995).

## Results

### Observation with intent to imitate

#### Comparing Deaf vs. Hearing groups

Overall sensorimotor EEG responses to ASL signs (both 1H and 2H stimuli combined) were not significantly different between Deaf and Hearing participants during the observation window. No significant differences were found using either full-scalp analyses or time-frequency visualizations at the electrodes overlying the sensorimotor ROI.

#### Sensitivity to sensorimotor characteristics

##### Time-frequency analyses across the scalp

For the Hearing group, there were no significant differences between conditions (1H and 2H) at any of the four time bins time during the observation window in any frequency band.

For the Deaf group, there were no significant differences between conditions in the lower alpha (8-10 Hz) or the upper beta (18-25 Hz) bands at any time. In the upper alpha (11-13 Hz) band, time bins 250-500ms, 500-750ms, and 750-1000ms showed more ERD in response to 2H signs compared to 1H signs. From 250-500 ms, the effect was seen a cluster of three electrodes (C2, CP2, and CP4) in the right central-parietal region. In the 500-750 ms bin, the effect was seen in a cluster of three electrodes (F6, AF8, and AF4) in the right frontal region. From 750-1000 ms, ten electrodes over the right posterior parietal region (CPz, CP2, CP4, TP8, P2, P4, P6, P8, POz, and PO4) showed significantly more alpha ERD in response to 2H signs.

Scalp activity in the lower beta (14-17 Hz) band also showed greater ERD in response to 2H signs in the Deaf group during the last three time bins. Topographic patterns of responses can be seen in Figure 2.

**Figure 2.**
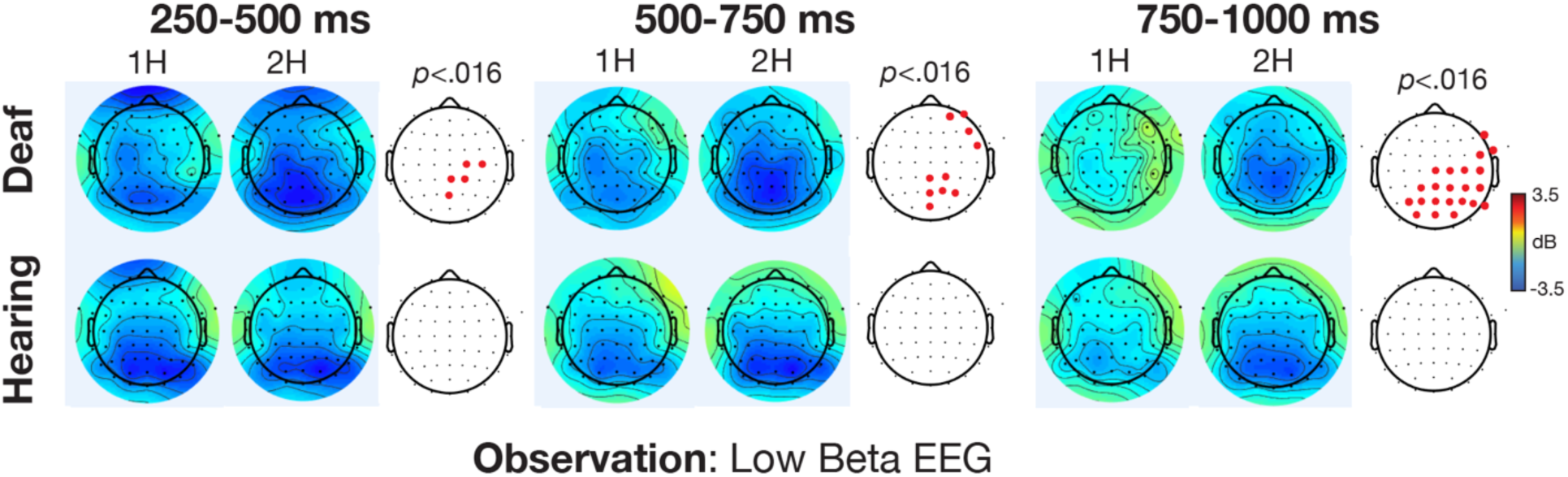
Low beta EEG power (14-17 Hz) during observation of signs with the intent to imitate, 250-1000 ms following onset of the sign in the video stimulus. Data are analyzed at 64 electrodes sites for Deaf and Hearing groups, in response to seeing one-handed (1H) and two-handed (2H) signs. Cool colors indicate desynchronization while warm colors show synchronization.

##### ROI analyses

We conducted targeted analyses at the electrodes overlying the central, fronto-central, and centro-parietal regions in order to assess the temporal dynamics of the responses to 1H and 2H signs while observing with the intent to imitate. Deaf and Hearing groups both showed greater ERD in response to 2H signs (*p* < .05, FDR corrected), although the profile of those responses showed a fair degree of variability. For example, at electrode CPz (see Figure 3), while both groups showed significantly more alpha and beta ERD in response to 2H signs, that difference was seen much earlier, and with more continuity, in the Deaf Signers compared to the Hearing Non-Signers. Eight electrodes within the ROI showed the same pattern of earlier and more robust discrimination of 1H vs. 2H conditions for Signers: C1, C2, C4, Cz, CP1, CP2, CP4, and CPz (pictured). In the other 13 electrodes in the ROI, the groups did not show a distinct difference in onset of discrimination between 1H and 2H signs. In all cases, for both groups, observation of 2H signs elicited greater alpha/beta ERD.

**Figure 3.**
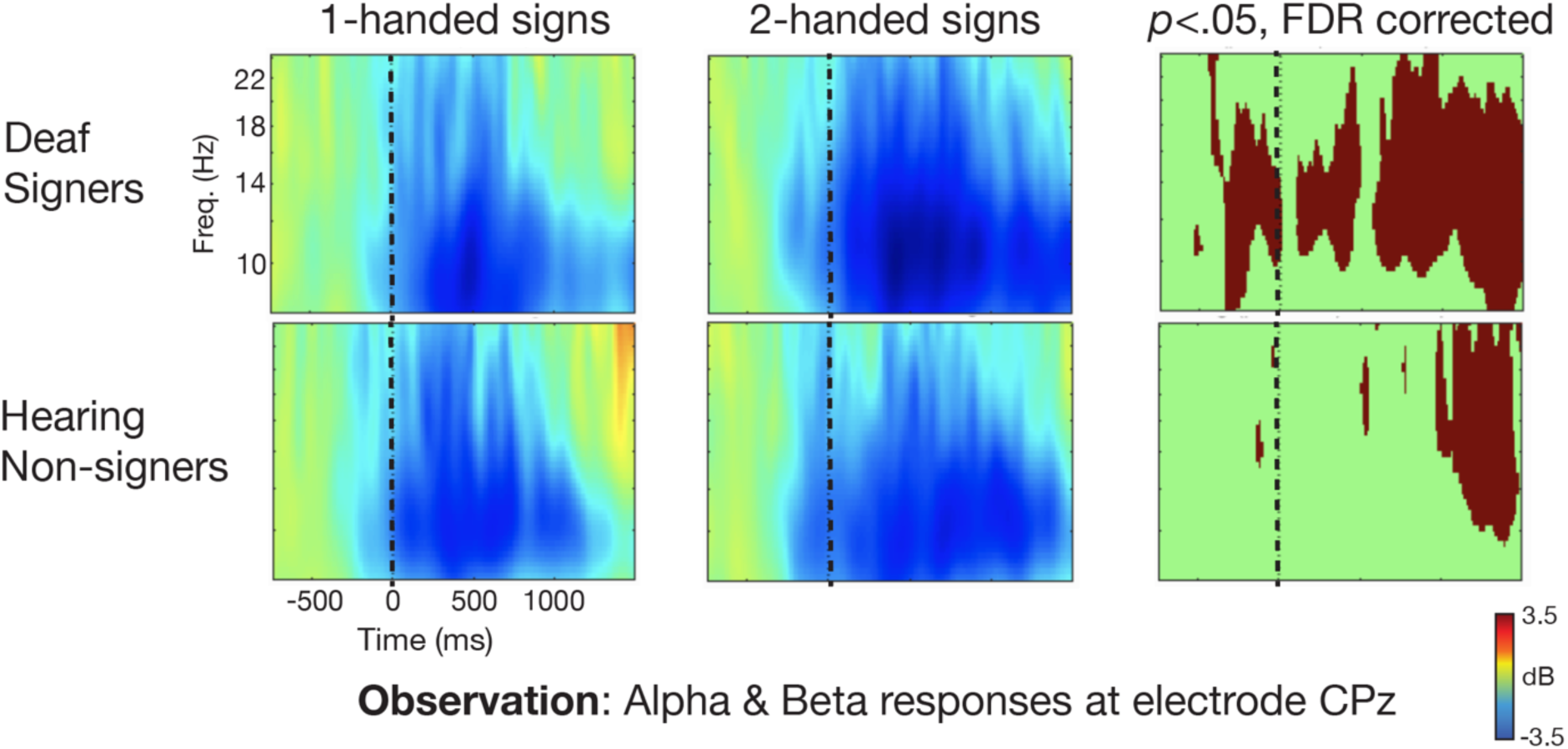
Comparison of alpha and beta activity during observation of signs with the intent to imitate, from -750 ms to 1500 ms (time 0 = onset of sign) at channel CPz for both groups. Deaf signers show earlier and more continual differentiation between 1H and 2H signs. Cool colors refer to event-related desynchronization relative to baseline.

### Producing ASL signs

#### Comparing Deaf and Hearing groups

##### Time-frequency analyses across the scalp

Deaf Signers and Hearing Non-signers showed no significant difference in alpha/beta EEG power during any of the first three production time bins (1000-1250 ms, 1250-1500 ms, or 1500-1750 ms). From 1750-2000 ms, at all four frequency bands, Deaf signers showed significantly lower EEG power compared to Hearing participants across a wide swath of electrodes including frontal, central, and right posterior scalp regions (see Figure 4 for visualization of the scalp distribution).

**Figure 4.**
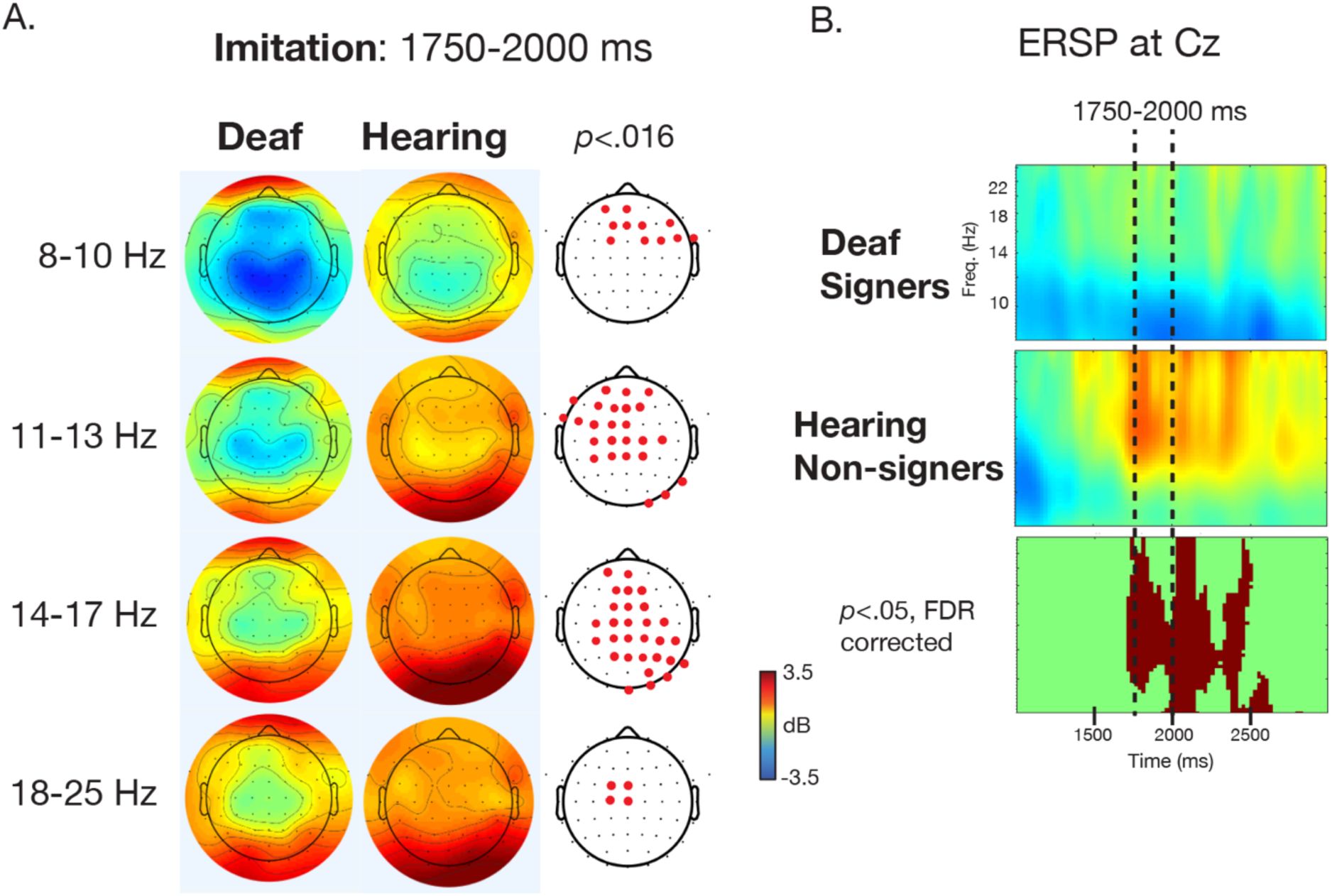
Alpha and beta frequency responses while participants imitate the signs. A) EEG responses across the scalp from 1750-2000 ms following the onset of the sign in the stimulus video. B) Time-frequency plot of activity at electrode Cz from 1000 to 3000 ms. The 1750-2000 ms period that is depicted in part A is marked with dotted lines.

##### ROI analysis

We conducted targeted analyses at the electrodes overlying the central, fronto-central, and centro-parietal regions in order to assess the temporal dynamics of the responses to all signs while imitating. The time-frequency analyses at the following eight electrodes showed significantly greater ERD in alpha/beta bands (p<.05, FDR corrected) in Deaf Signers compared to in Hearing Non-Signers: FC1, FCz, C1, C2, Cz, CP1, CP2, and CP6 (see Figure 4B for one representative plot).

#### Sensitivity to sensorimotor characteristics

##### Time-frequency analyses across the scalp

In the lower alpha (8-10 Hz) band, Hearing Non-Signers showed stronger alpha ERD at five right parieto-occipital electrodes (P4, P6, P8, PO8, O2) when producing a 2H sign during the 1000-1250 ms time bin. No other differences were seen at any time for the Deaf or Hearing groups in the low alpha range.

In the upper alpha (11-13 Hz) band, both the Deaf and Hearing groups showed significantly more alpha ERD as they produced 2H signs during all four time bins. See Figure 5 for the topographical distribution of these effects across the scalp across the latter three time bins.

**Figure 5.**
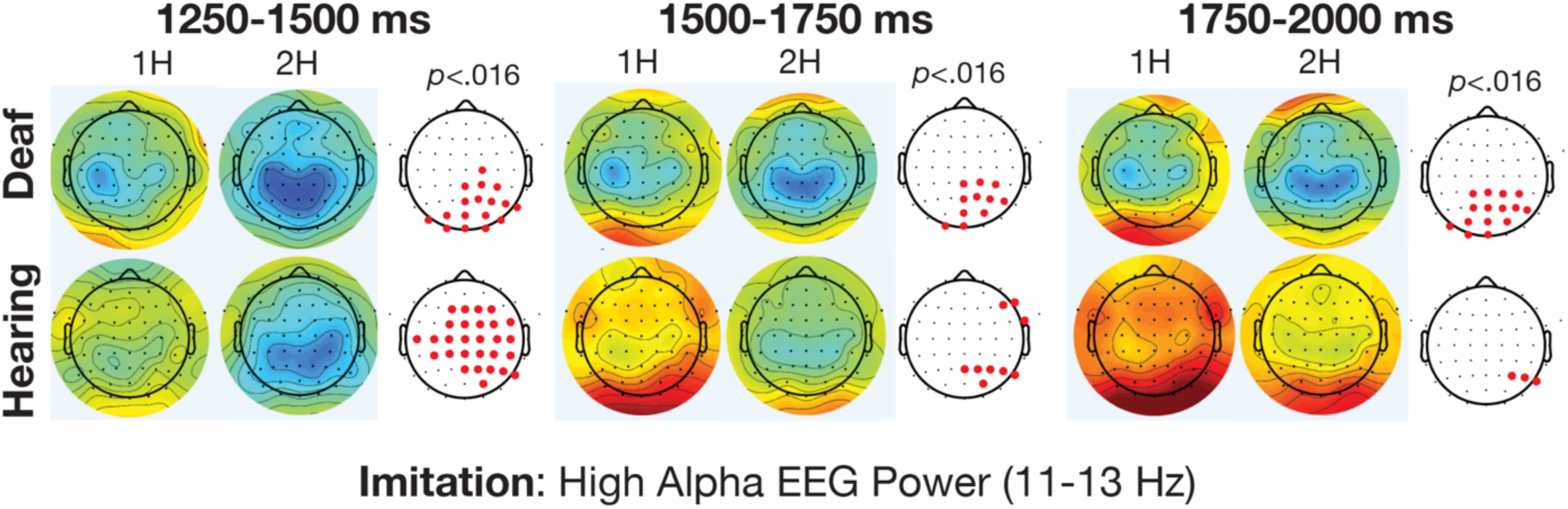
High alpha EEG power (11-13 Hz) while participants imitate one-handed (1H) and two-handed (2H) signs, from 1250-2000 ms following onset of the sign in the video stimulus. Data are analyzed at 64 electrodes sites for Deaf and Hearing groups, in response to seeing one-handed (1H) and two-handed (2H) signs. Cool colors indicate desynchronization while warm colors show synchronization.

In the lower beta (14-17 Hz) band, both the Deaf and Hearing groups showed differentiation while producing 1H and 2H signs. These effects were present across vast regions of the scalp more beta ERD during 2H sign production compared to 1H sign production across fronto-central, central, parietal, and occipital regions from 1000-1250 and 1250-1500 ms. During these time bins, effects were present for both groups at more than 50% of the electrodes, spanning broad regions of the scalp. From 1500-1750 ms, the differences became less widespread and were apparent at seven parietal electrodes in the Deaf group and 17 right frontal and parietal electrodes for the Hearing group. From 1750-2000 ms, there were no significant differences for the Hearing group, while the Deaf group showed more beta ERD during production of 2H signs at 14 bilateral centro-parietal electrodes.

In the upper beta band (18-25 Hz), both groups displayed greater beta ERD during production of 2H signs at centro-parietal electrodes from 1000-1250 ms. For the hearing group this effect was seen across 18 bilateral central, parietal, and occipital electrodes, whereas for the Deaf group the effect was present at 6 right parietal electrodes only. From 1250-1500 and 1500-1750 ms, the Hearing group showed significantly lower beta power for 2H signs at more than 50% of scalp electrodes, and the Deaf group showed no differences. There were no beta ERD differences in the last time bin.

##### ROI analysis

The targeted ROI analyses revealed that both Deaf and Hearing groups showed significant differences (*p* < .05, FDR corrected) in EEG activity between conditions over the central region during sign imitation. However, these differences came about as a result of starkly different profiles of activity in the sensorimotor cortices for the two groups. For the Hearing group, much of the difference was driven by increases in EEG power across alpha and beta ranges, particularly when producing a 1H sign. For example, at electrode CPz (see Figure 6), while both groups showed significant differences in power between 1H and 2H signs, for the Deaf group that effect is driven by alpha/beta ERD, with more ERD in response to 2H signs. In contrast, for the Hearing group, both conditions elicit an increase in EEG power, with higher power for 1H signs and lower power for 2H signs. For both groups these differences were apparent throughout the production window. These patterns were widespread across the ROI, with significant effects occurring in the same direction at all electrodes in the ROI for the Hearing group, and for 18 electrodes in the ROI for the Deaf group.

**Figure 6.**
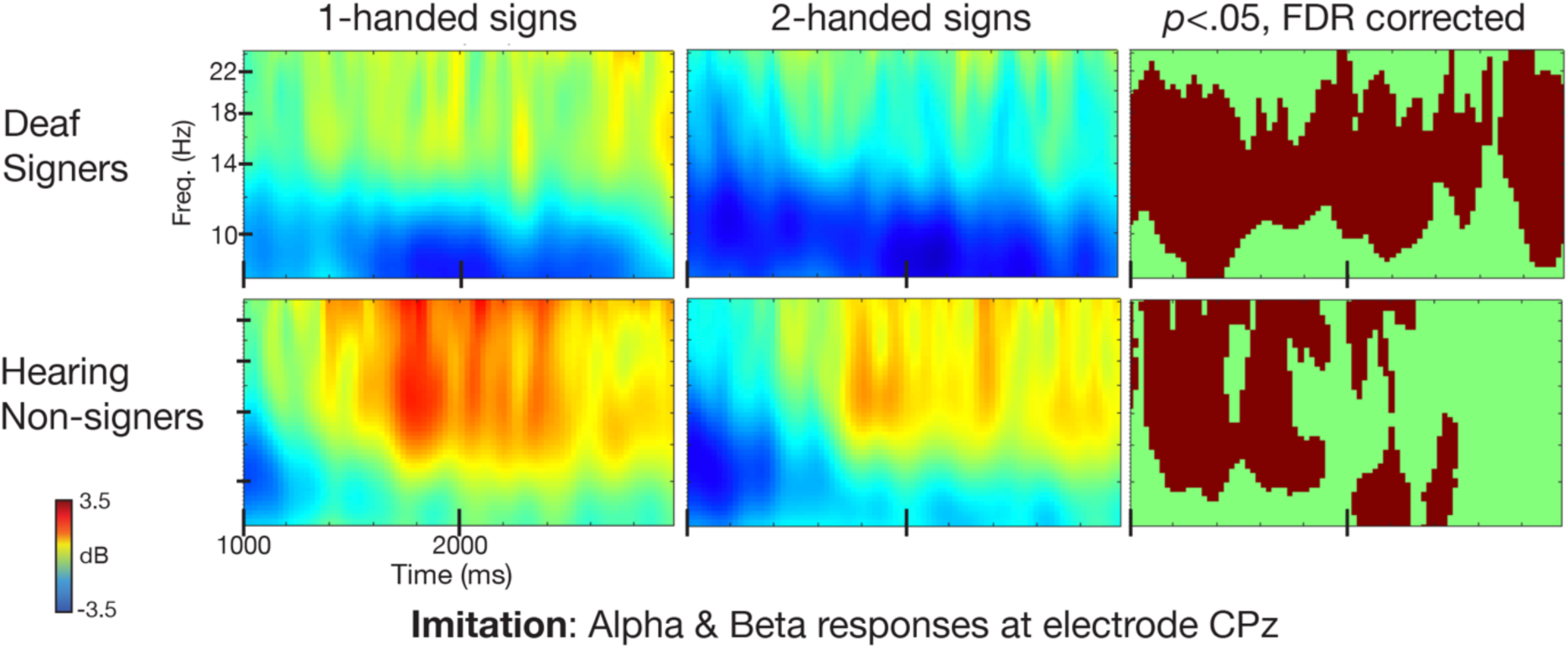
Comparison of alpha and beta activity during imitation, from 1000 to 3000 ms (time 0 = onset of sign) at channel CPz for both groups. Cool colors refer to event-related desynchronization relative to baseline.

## Discussion

In the current study, we questioned how experience with American Sign Language impacts sensorimotor EEG activity during an imitative signing task. This question was motivated by interests in how action experience influences the neurodynamics of action processing, and also by open questions about the sensorimotor processing of signed languages. We analyzed sensorimotor EEG activity in alpha and beta frequency bands while participants watched and imitated individual ASL signs. We analyzed data both from the period of time when participants were watching with the intent to imitate, and during the period of time when they were carrying out their imitation.

### Watching with the intent to imitate

We predicted that during sign observation, Deaf signers would show less sensorimotor system activity and less differentiation of sign types in the sensorimotor system, because of the possibility that for those individuals the task would involve language systems of the brain more robustly than sensorimotor systems. This hypothesis was based on prior work using a similar paradigm in a passive (non-imitative) task (Kubicek & Quandt, 2019), as well as other functional neuroimaging evidence (Corina & Knapp, 2006; Emmorey, Xu, Gannon, Goldin-Meadow, & Braun, 2010; Okada et al., 2016; Rogalsky et al., 2013). However, in opposition to our prediction, we found no significant differences in overall alpha/beta ERD between the two groups while participants were observing the signs. This is likely due to the imitative context of the current study. Given that that participants were all watching signs with an intent to imitate them, the typical neural response seen during action observation was likely overlaid with the neural substrates of motor preparation. This preparatory anticipation may have engaged Deaf Signers’ sensorimotor systems more readily. Recent work comparing action experts and novices has shown greater alpha ERD when the experts anticipate upcoming actions, an effect seen over frontal, occipital, and parietal brain regions (Simonet et al., 2019). It is possible that the context of motor preparation in our task increased the signers’ alpha/beta ERD due to their greater expertise with the signs, thus washing out any significant differences between the groups that would be seen in a non-imitative paradigm.

We expected that both groups would show different sensorimotor EEG responses to one-handed and two-handed signs, but that the effect would be stronger in the Hearing Non-Signers, due to their unfamiliarity with the signs, lack of conceptual understanding of their meanings, and the resultant need to focus on the basic physical parameters of the observed gesture. However, our data did not support this prediction. The Hearing group showed no significant differences between conditions (1H and 2H) during the observation window in any frequency band. For these individuals, sensorimotor differentiation between conditions did not start until after our observation window was over. However, deaf signers showed earlier and more consistent differentiation between one- and two-handed signs. Two-handed signs elicited greater alpha and beta ERD in Deaf Signers, as seen by the cluster of right-central electrodes which differentiated between stimulus types during the 250-500 ms time bin (see Figure 2). Taking a closer look at the temporal dynamics of neural oscillatory activity in the central region, we see that Deaf Signers’ sensorimotor response differentiates extremely early between 1H and 2H signs—before the actual onset of the sign. Indeed, as seen at electrode CPz (in Figure 4), there is significantly greater ERD for two-handed signs starting ∼300 ms before sign onset, while the model’s hands are moving into place to begin producing the sign. Thus, we show that fluent deaf ASL users engage mirroring-like processes earlier, and more continuously, than do hearing non-signers when seeing ASL signs with an intent to imitate them.

The deaf signers in our study showed discrimination between observed one- and two-handed signs very early. Our findings complement prior work demonstrating that deaf signers discriminate between plausible and non-plausible signs quickly, within around 100 ms of seeing a still image representing a sign (Almeida et al., 2016). The timing of the effects we see in our study aligns with prior work as well. In one action production task, participants’ beta ERD was seen within 110-120 ms of seeing a cue about the upcoming action (Tzagarakis et al., 2010). Our results suggest that sign language experience can tune action mirroring processes in the brain to more quickly discriminate between the specific sensorimotor characteristics of observed signs.

In many ways, our results echo prior findings from work looking at how action expertise changes the mirroring-related processing of others’ actions. Recent work comparing action experts and novices sheds light upon the current findings. For instance, high alpha ERD is associated with the activation of specific sensorimotor characteristics of observed actions (Denis et al., 2017), likely because experts have greater access to calling upon the specific sensorimotor characteristics of an observed action than do non-experts. While our analyses did not reveal significant group differences in high alpha ERD during observation, Signers did show significantly more sensitivity to the specific sensorimotor characteristics of actions in the high alpha band, compared to Non-Signers. Specifically, our signing group showed early high-alpha and low-beta ERD in response to two-handed signs, compared to one-handed signs. This sensitivity to the motor characteristics of the observed action in these frequency bands suggests that signers are calling upon their prior sensorimotor experiences with the signs they are seeing, and invoking mirroring-like simulation as they see the action unfold.

In other domains of human action (e.g., dance, grasping), the action experts’ sensorimotor cortex is more active while they see others perform actions with which they have experience, like we see in the current results. However, in the this study, there is a vast discrepancy between the amount of ASL experience that the Deaf Signers and Hearing Non-Signers have, and if we were able to compare sensorimotor reactivity across the full spectrum of ASL experience (e.g., including hearing fluent signers, and intermediate signers), it is very possible that a more complex, non-linear relationship between sign experience and mirroring would emerge (Gardner et al. 2017a, 2017b; Gardner et al., 2015).

Some of the key findings in this paper involve greater neural activity in response to two-handed signs over the right sensorimotor cortex (i.e., right central/centro-parietal electrodes). Given that all our signers were right-handed, and the model was right-handed, this pattern suggests that participants were indeed simulating the production of action as they observed, or perhaps invoking preparatory motor plans in anticipation of imitation. One-handed signs in ASL are always produced with the dominant hand, which is associated with primariliy contralateral sensorimotor cortex activity. Two handed signs, in contrast, invoke bilateral sensorimotor cortex (Emmorey et al., 2016). Thus, the predicted difference between two-handed and one-handed signs, if an observer is calling upon their own sensorimotor cortices to process observed actions, would be over the right-lateralized sensorimotor cortex. This is what we found in high alpha and low beta EEG rhythms during the observation window, particularly in early time bins (e.g., 250-500 ms) for Deaf Signers only.

It is likely that in the current study, those effects were heightened due to the imitative context in which the signs were observed. The task of preparing to imitate likely primed observers (both Hearing and Deaf) to prepare their own motor plans for reproducing the signs. Watching sign language has been associated with the generation of internal predicting coding models, as indicated by increased activity in the superior parietal cortex during sign viewing compared to speech listening in hearing bimodal bilinguals (Emmorey et al., 2014). Our current results suggest that indeed, fluent signers are encoding the articulatory specifications of observed signs from a very early time during sign viewing, a phenomenon which appears to be enhanced in an imitative context, where internal predictive models supporting comprehension may be working in parallel to motor planning in advance of producing one’s own imitation.

### Producing ASL signs

In the current paradigm, participants experienced a predictable imitative exchange. In each trial, they saw a sign, then produced the sign themselves. This consistency means that in this context there was no strict boundary between sign observation and when participants were preparing their own imitations. However, we opted to separately analyze the time while participants were initiating and carrying out their own productions of the signs. Our goal was to see whether mirroring systems would engage more greatly during this period for the expert signers, who have a great deal of sensorimotor experience to draw upon, but who may be processing the signs largely linguistically, or for the hearing novices, for whom the signs represent complex gestures they likely have never produced before.

There were stark differences in EEG activity during production between the two groups. Overall, the Deaf group showed a sustained ERD response across alpha and beta frequencies, whereas the Hearing group showed increased power across these frequency bands (see Figure 4). This difference between the two groups during action production was in contrast to our expectations. We had expected that while imitating signs, Hearing Non-Signers would recruit sensorimotor regions more than Deaf Signers, as in our prior work wherein Hearing Non-Signers showed greater alpha/beta ERD when passively observing signs, in comparison to Deaf Signers (Kubicek & Quandt, 2019). However, what we found suggests that instead, during the actual production of ASL signs, Deaf Signers recruit sensorimotor cortex in a typical way, exhibiting ERD over central sites, predictably differentiating between one- and two-handed signs. In contrast, Hearing Non-Signers show less ERD over central sites, and in fact show increases in alpha and beta EEG power as they carry out actions.

The observed pattern of results suggests that in this imitative paradigm, Deaf Signers are involving their sensorimotor cortices as they produce signs they are undoubtedly very familiar with. This suggests that although the Deaf Signers know the semantic and other linguistic features of the signs, and have a lifetime of experience both perceiving and producing the signs, their sensorimotor cortices are still underlying their sign productions. This should come as no surprise, given that signed communication requires coordinated movements of the fingers, hands, arms, and body. As well, our results align with prior PET findings of signers producing ASL signs (Emmorey et al., 2016). Both comprehending and producing ASL signs preferentially engages the premotor cortex, parietal cortices, and motion-sensitive areas of the middle temporal gyrus moreso than when speaking or comprehending speech (Emmorey et al., 2014). This pattern is likely due to the gross sensorimotor demands of coordinating and articulating language using the hands and body, and the demands of perceiving another person’s complex hand and body articulations. Looking to the broader action-expertise literature, our current results speak to the notion that in some circumstances, action experts do exhibit greater involvement sensorimotor cortices during action than do action novices, in contrast to the idea of neural efficiency (Babiloni et al., 2010; Babiloni et al., 2009).

The specific oscillatory activity within the four frequency bands we studied can yield further information about what characteristics of action were contributing to these effects. During sign production, low alpha power at central region electrodes did not differ between groups (although frontal regions differed), perhaps because the low alpha rhythm is thought to represent broad initiation of movement, without much differentiation between movement types (Denis et al., 2017; Pfurtscheller & Neuper, 2000). Greater ERD within the beta frequency band is thought to reflect a higher degree of certainty about how to produce an action (Palmer, Zapparoli, & Kilner, 2016; Tzagarakis et al., 2010). In comparing beta ERD between the two groups (regardless of 1H/2H conditions) during the production window, hearing non-signers exhibited a surge of higher power in the beta frequency band, which may be because of their uncertainty about how to proceed. In contrast, the deaf signers showed a clear beta ERD response, perhaps due to their certainty and existing knowledge about how to produce the sign they had just seen.

When comparing EEG activity between 1H and 2H conditions, Deaf Signers showed a predictable result: greater sensorimotor cortex activity (as shown by more alpha/beta ERD) while producing a 2H sign. As expected, this effect was evident over right centro-parietal electrodes, suggesting that the right sensorimotor cortex is more involved when carrying out a movement that includes the left hand and arm. However, the Hearing Non-Signers displayed an unexpected pattern of results. While EEG activity was significantly lower in the 2H condition, there were overall increases in EEG power for both conditions, and the difference appears to be driven by a marked increase in EEG power during the production of 1H signs (Figures 5 and 6).

### Hearing non-signers imitating signs

In our task, the hearing non-signers likely experienced a great deal of uncertainty, given the challenge of reproducing novel signs after only one viewing. The difficulty of the task for Hearing Non-Signers likely had a strong influence over these unexpected results. For hearing non-signers, learning ASL signs draws upon working memory capacity, particularly phonological short-term memory (Martinez & Singleton, 2018). Beta event-related synchronization at frontal electrode sites is greater during simple working memory tasks compared to a more difficult task (Pesonen, Hämäläinen, & Krause, 2007). Recent work also suggests that beta activity (14-20 Hz) in the parietal cortex serves as an episodic buffer, holding recent sensory input and linking it to executive commands to produce relevant actions (Gelastopoulos, Whittington, & Kopell, 2019). It is possible that for Hearing Non-Signers, there was a difference in the degree of working memory and episodic buffering involved for the one-handed and two-handed signs, which could help explain the effects seen in that group.

The hearing non-signers in our experiment may have been stressed by the task at hand, prompting them to develop various strategies to hold the motor plans in place before imitation occurred, and necessitating mental rotation-related skills due to the need to produce an action which had only been seen from the third-person perspective (Shield & Meier, 2018). Hearing participants’ development, testing, and use of these various strategies (e.g., covert rehearsal, attempts to make semantic links, attempting to memorize motor sequences) may be a significant source of mental effort in a task such as ours (Martinez & Singleton, 2018), which likely influences the neural oscillatory patterns during sign production (Gelastopoulos et al., 2019). This aligns with other prior research showing that hearing non-signers show increased effortful attention during sign perception (Williams, Darcy, & Newman, 2016). The difficulty of the task for the hearing group stands in contrast to its ease for the Deaf Signers, who were essentially doing a word-shadowing task using words they were familiar with.

### Future Directions

While the results presented here contribute to our growing understanding of how deaf signers and hearing non-signers process visual stimuli differently, there are many questions left unanswered. One limitation of this study is the lack of different expertise levels with ASL, which would allow us to observe how the neural responses to perceiving and producing ASL change with expertise. Our group of Deaf Signers was quite heterogeneous—the group included native signers who grew up using ASL, as well as individuals who were fluent in ASL after learning it later in life. This heterogeneity provides strength to our findings, in that the effects were present even in such a diverse group, but it limits the ability to specify whether the observed differences are due more to deafness, or to fluency with ASL. Future inclusion of other groups (e.g., hearing native ASL users, or deaf people who don’t know sign language) would significantly clarify this issue. More narrow recruitment criteria would likely yield more detailed information about how ASL perception in native signers may differ from those who became fluent later in life (Twomey, Price, Waters, & MacSweeney, 2019). Finally, the results we present here cannot differentiate precisely between the sensory plasticity arising from deafness and the effect of fluency in ASL. It is our hope that future work can further disentangle these complex, and likely overlapping, effects.

## Conclusion

Fluent deaf sign language users have long-term experience with producing and perceiving the complex gestures that make up signed languages. To date, the research about whether, and how, sign language users may invoke the mirroring system during sign perception has yielded mixed results. We conducted this EEG study to assess the timing and presence of mirroring-like processes during the perception of American Sign Language signs, and the profile of sensorimotor involvement during the imitation of the observed signs. We present evidence that deaf ASL signers show earlier and more robust involvement of their own sensorimotor cortices to discriminate the sensorimotor characteristics of observed signs, in contrast to hearing non-signers. When producing their own versions of the observed signs, stark differences were apparent in the oscillatory responses across all measured EEG frequency bands. Together, this work demonstrates that in an imitative context, ASL users rapidly process others’ signs by drawing upon their own sensorimotor representations of those signs. It appears that fluent deaf signers are particularly sensitive to the physical characteristics of observed signs, which is yet another way in which deaf signers show enhanced motion perception.

## Acknowledgements

We are grateful to all participants who gave their time for this study. We recognize the support of Emily Kubicek, Naseem Majrud, and Taylor Wardle in collecting this data. This work was supported by the Ph.D. in Educational Neuroscience program at Gallaudet University and by National Science Foundation grant #1839379 to LQ. The authors declare no conflict of interest regarding this work.

